# Systematic assessment of homology-based methods for fine-grained functional annotation using diverse protein families

**DOI:** 10.1101/2025.09.17.676781

**Authors:** Rakesh Busi, Pranav Machingal, Nandyala Hemachandra, Petety V. Balaji

## Abstract

The size of the protein sequence database is increasing without a consequent increase in the number of proteins with known molecular function, especially at the fine-grained level. Alignment-based approaches such as BLAST and profile hidden Markov models (HMMs) are widely used to infer homology and transfer annotation to be subsequently confirmed by experiments. The ability of BLASTp to distinguish orthologs from paralogs varies across protein families; for profile HMMs, this depends on the sequences considered for generating multiple sequence alignments. In this study, we systematically evaluated the performance of BLASTp and HMM-based methods for fine-grained function annotation using carefully curated protein datasets that are diverse in sequence-structure-function relationships. Expectedly, BLASTp performed well in detecting close homologs but failed to detect remote homologs. BLASTp detected homology between 22.6% and 100% of sequence pairs within different homologous protein families. The extent of sequence identity between trypsin and chymotrypsin sequences is high despite differences in fine-grained molecular function. Transferring function annotation based on homology inferred from BLASTp leads to errors in trypsin-chymotrypsin-like situations.

Profile HMMs improved sensitivity and captured subtle homology signals even when sequence identity was low, though some known family members scored below threshold due to functional divergence or mutations at catalytic sites. We further showed that relying solely on homology for annotation transfer can lead to misleading conclusions when proteins have evolved divergent functions despite structural similarity. Our findings highlight that a cautious approach involving BLASTp, profile HMMs, and expert domain knowledge provides the most reliable strategy for functional annotation. After all, not every family member may be doing what we think they are doing.

## Introduction

Gene or genome duplication followed by functional divergence leads to genes encoding new functions. Several models have been proposed to explain functional divergence during evolution. These include (i) neofunc-tionalisation, (ii) subfunctionalisation by duplication, degeneration, and complementation, (iii) subfunction-alisation via specialisation or escape from adaptive conflict, and (iv) innovation-amplification-divergence [1]. With reference to the function of the ’parent gene’, the new function may be a variant or completely new. Trypsin and chymotrypsin are examples of the former [2], [3], and lysozyme c and *α*-lactalbumin are examples of the latter [4].

Orthologs arise from speciation events and typically retain the same function in different organisms, showing high sequence identity when the species are closely related. Paralogs, in contrast, originate through duplication within a genome and may diverge in function. Both orthologs and paralogs are homologs, sharing sequence and structural similarities [5]. However, such similarities often lead to uncertainties in distinguishing paralogs and orthologs from each other.

The BLASTp algorithm can be used to search for homologs of a query sequence in a database [6]. Homology is inferred if (i) a high percentage query sequence is aligned and (ii) sequence identity ≥30% [7]. Several homologs show high sequence divergence, and sequence identity between such pairs will be <30% ^1^. However, such proteins share the same fold. Homologs that show only fold similarity and low sequence identity are considered remote homologs. It is important to note that BLASTp misses remote homologs due to their low sequence identity. Therefore, two sequences that are not aligned by BLASTp are either remote homologs or not homologs.

BLASTp is inherently ”blind” to biological functions; it does not assess functional relationships but rather detects evolutionary relationships based on sequence identity. Hence, it is not possible to know whether the hit sequence is an ortholog or a paralog of the query unless additional information is available. Sequence alignments are often not end-to-end because of high sequence divergence (remote homologs), truncations, indels, and/or gene fusions. These factors limit the use of BLASTp for function annotation based on sequence similarity.

Profile methods, e.g., PSI-BLAST [10] and profile HMM [11], exploit the availability of sequences of several homologs to identify remote homologs with higher confidence relative to what one can achieve by BLASTp. These methods are critically dependent on two factors: (i) ensuring that only homologs are used for building profiles and (ii) the quality of the multiple sequence alignment (MSA). If one were to use the profiles for function annotation, then it becomes important to use only structural and functional homologs for building a profile. Functional homology is essential to avoid inclusion of paralogs. As for the quality of the MSA, currently, there is no widely accepted metric other than visual analysis of sequence conservation and gap positions.

BLASTp is being used extensively to detect sequence similarity since its introduction about 3 decades ago. And sequence similarity is used to infer homology and transfer of molecular function annotation. Several studies have shown that homologs can be quite divergent (remote homologs; inference based on fold similarity) and, in many cases, even the function diverges (paralogs). The focus of this study is fine-grained function annotation, i.e., the 4th digit of the EC number. The objective of the present study is to systematically assess if fine-grained molecular function can be inferred based on homology detected by BLASTp and profile HMM. Towards this objective, we have considered 15 datasets, and each dataset consists of two or more protein families. These datasets are divided into 5 groups on the basis of the fold-function relationship. Similarities and differences in fold and function between the protein families show wide variations (Table 1). In all, 5224 single domain sequences representing 26 protein families and protein sets like non-SNARE, non-GPCR (TM), and non-GPCR (no TM) were considered.

**Table 1.** Grouping of datasets based on fold and function relationships.

| Grp. | Datasets | Fold and function relationships between the families <sup>$\phi$</sup> | Remarks about protein families within a dataset |
| --- | --- | --- | --- |
| 1 | Lysozyme CGCh (3 families) | Fold - Different<br>Function - Similar | (i) Not homologs since no fold similarity.<br>(ii) Convergent evolution since functional similarity of non-homologs. |
|  | Xylanase (2 families) |  |  |
|  | Chitinase (2 families) |  |  |
| 2 | Lysozyme CaLA (2 families) | Fold - Same<br>Function - Dissimilar | (i) Homologs since fold similarity.<br>(ii) Paralogs since no functional similarity. |
|  | Protease (2 families) |  |  |
|  | Ferredoxin (2 families) |  |  |
| 3 | β-Glucosidase (2 families) | Fold - Same<br>Function - Similar | (i) Homologs since fold similarity.<br>(ii) Orthologs since functional similarity. |
|  | Lysozyme (3 families) |  |  |
| | β-Galactosidase (3 families) | | |
| 4 | GH2 (2 families) | Fold - Same<br>Function - Dissimilar | (i) Homologs since fold similarity.<br>(ii) Paralogs since no functional similarity. |
|  | GH3 (2 families) |  |  |
|  | GH5 (2 families) |  |  |
| 5 | SNARE (2 families) | Fold - Different<br>Function - Dissimilar | Completely unrelated both in terms of function and evolution. |
|  | GPCR1 (2 families) |  |  |
|  | GPCR2 (2 families) |  |  |

The table summarizes the grouping of 15 datasets into 5 groups according to different combinations of fold and function relationships between protein families. It also provides remarks on the presence of homologs, paralogs, and orthologs within each dataset. Notably, Group 2 and Group 4 share the same fold and function relationship, while Group 4 uniquely contains GH (Glycoside Hydrolase) labels.
<sup>$\phi$</sup> Based on literature reports.

In Table 1, the datasets with the same fold are considered homologous [12] and datasets with paralogous protein families, i.e., same fold but dissimilar function, were grouped into Group 2 and Group 4, with Group 4 exclusively containing GH-labelled sequences. The datasets in Group 3 contain orthologous families that share the same fold and have similar functions. In contrast, the datasets in Group 1 are not homologous, as they differ in fold and are likely functional analogs arising from convergent evolution. The datasets in Group 5 contain completely unrelated protein sets.

## Methods

### Datasets

The choice of protein pairs for comparison was based on (i) domain knowledge and (ii) the number of sequences available. This led to the datasets Xylanase, Chitinase, *β*-Glucosidase, Lysozyme, *β*-Galactoside, GH2, GH3, and GH5 (Table 1). These constitute Groups 1, 3, and 4. Lysozyme CGCh was specifically considered because of the availability of extensive experimental (ground truth) data. Here, ”extensive” means that we can find at least one literature reference (PubMed ID) to every sequence. Notably, some of the sequences are present in more than one Group (Table 2).

**Table 2.** Summary of sequences present in multiple Groups.

| Number of sequences | Group where the sequences are represented |  |
| --- | --- | --- |
| 26 | Protein family: Lysozyme C<br>Dataset: Lysozyme CGCh<br>Group: 1 | Protein family: GH22<br>Dataset: Lysozyme<br>Group: 3 |
| 16 | Protein family: Lysozyme G<br>Dataset: Lysozyme CGCh<br>Group: 1 | Protein family: GH23<br>Dataset: Lysozyme<br>Group: 3 |
| 117 | Protein family: GH2<br>Dataset: β-Galactosidases<br>Group: 3 | Protein family: β-Galactosidases<br>Dataset: GH2<br>Group: 4 |
| 204 | Protein family: GH3<br>Dataset: β-Glucosidase<br>Group: 3 | Protein family: β-Glucosidase<br>Dataset: GH3<br>Group: 4 |

The table summarizes sequences that are present in multiple Groups with different protein family labels. It shows the number of such sequences and details the protein families, datasets, and Groups in which these sequences are commonly found.

Protein pairs within any given dataset belonging to Groups 1 to 4 share similarities in at least one of these two characteristics: fold and function. Group 5 datasets were chosen as ’control’ wherein the comparison is between two sets: one set consists of homologs that have the same coarse-grained annotation, and the second set was chosen such that sequences do not share fold or function similarity. This group consists of SNAREs and GPCRs.

The SNARE family proteins were chosen based on coarse-grained function annotation [13]. Proteins were considered without taking into cognizance the category to which they belong, i.e., v-SNARE, t-SNARE, R-SNARE, or Q-SNARE. This suggests that the SNARE family of proteins show a high degree of sequence divergence, and this is in consonance with (i) choosing SNARE proteins irrespective of the category to which they belong and (ii) the large variations in the sequence lengths.

The only commonality to all GPCRs sequences considered in this study is the 7-transmembrane (7-TM) domain architecture. Non-GPCRs were chosen in two ways: (i) they have at least one TM domain (non-GPCR™) and (ii) they should not have even one TM domain (non-GPCR(no TM)). Clearly, in Group 5, comparison is between diverse sets of proteins, viz., GPCRs or not, and have/do not have SNARE activity.

### Fetching and pre-processing of sequences

The GenBank IDs for the protein families present in the datasets of Groups 1, 3, and 4 were obtained from the CAZy database, while the corresponding sequences were retrieved from Batch Entrez of NCBI. For the protein families in the Protease dataset, the UniProt IDs were taken from Brenda [14], and the corresponding sequences were downloaded from UniProt [15]. The sequences for *α*-Lactalbumin, SNARE, and non-SNARE protein families were obtained from relevant literature. For all the remaining protein families, both the IDs and sequences were directly fetched from UniProt (Table 3).

**Table 3.**
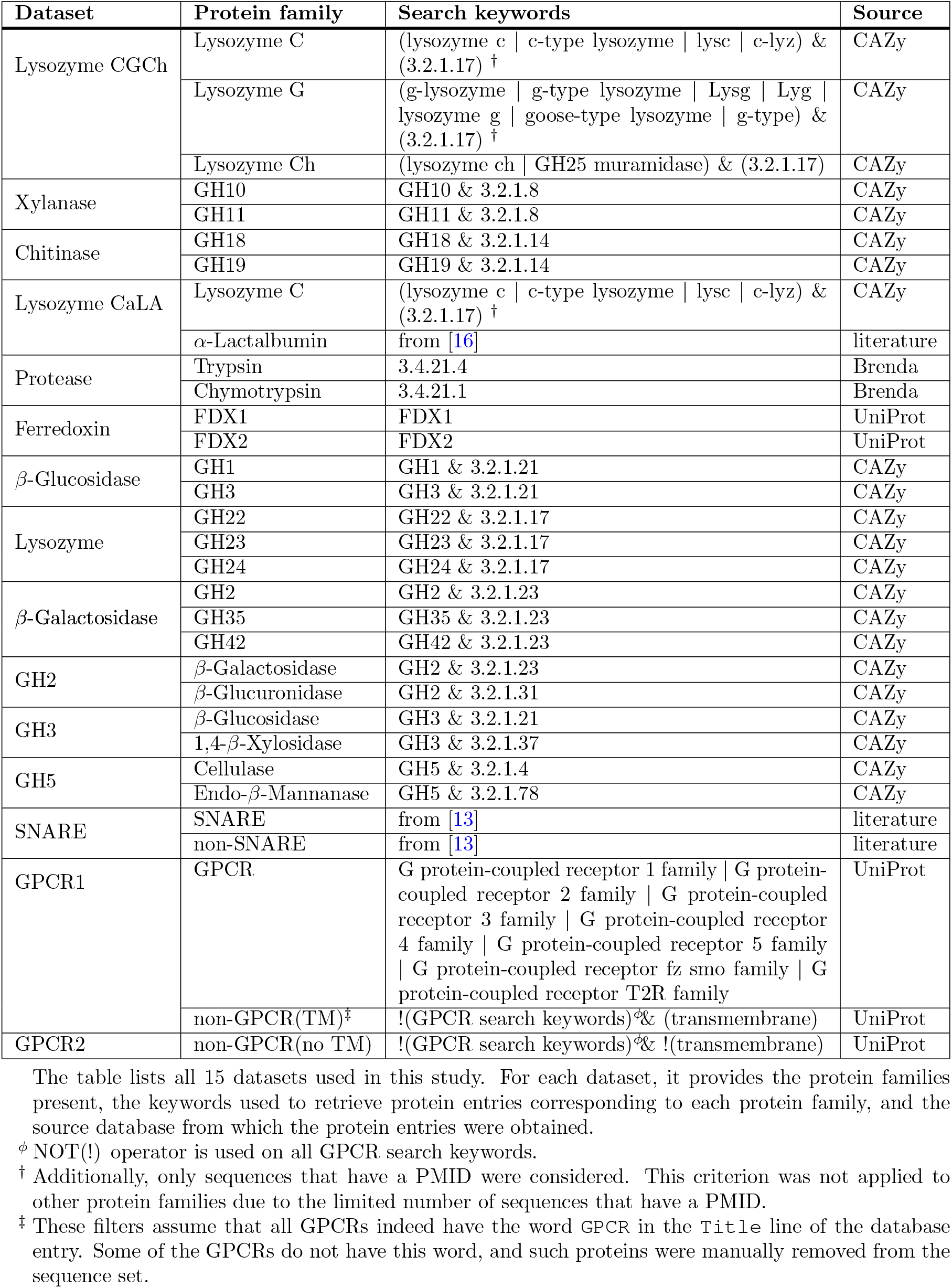
Details of datasets, protein families, search keywords, and sources.

The fetched sequences were subjected to the following preprocessing steps: (i) sequences with headers containing either the term `partial` or `fragment` were removed because we wanted to consider only full length sequences, (ii) sequences with two or more EC numbers were omitted because we did not want to include fusion proteins with ≥2 domains and protein which have more that one activity associated with the same active site, (iii) sequences with duplicate accession numbers between protein families were removed, while within each protein family, only a single copy of entries with duplicate accession numbers or identical sequences was retained, and (iv) sequences containing the letters B, J, O, U, X, and Z were removed, as these letters do not represent any of the 20 standard amino acids. Additionally, outlier sequences, if any, were removed based on the sequence length distribution of the protein family, as determined by a box-whisker plot.

### Pairwise sequence alignment

All-against-all pairwise sequence alignment was performed to determine the percentage sequence identity (PSI) and percentage query coverage (PQC) for all protein sequence pairs of our datasets. NCBI’s Protein-Protein BLAST 2.9.0+ (BLASTp) command-line application was executed on a Linux server using default values for all the parameters unless mentioned to the contrary. The command executed is `blastp -query dataset.fasta -subject dataset.fasta -outfmt “6 qseqid sseqid pident qcovhsp” -out result.txt.` Here, both the -query and -subject parameters were set to the same file `dataset.fasta` to enable all-against-all pairwise comparison within a dataset. The output `result.txt` was generated in tabular format `outfmt 6` with four columns: `qseqid` (query sequence ID), `sseqid` (subject sequence ID), `pident` (percentage of identical matches), and `qcovhsp` (query coverage per high-scoring segment pair (HSP)). An HSP refers to a local, gap-free alignment with one of the highest scores in the search. For bidirectional hits or multiple alignments of the same pair, we retained the result with the highest `qcovhsp` (maximizing alignment length). Two sequences were inferred to be homologs if the following criteria are met: (i) percentage sequence identity threshold (*T*_%ID_) ≥30% [7]; also used in AlphaFold [17] and Seq2Symm [18] and (ii) percentage query coverage threshold (*T*_%QC_) ≥70% (Fig 1 in Supporting information S1 File).

**Fig 1.**
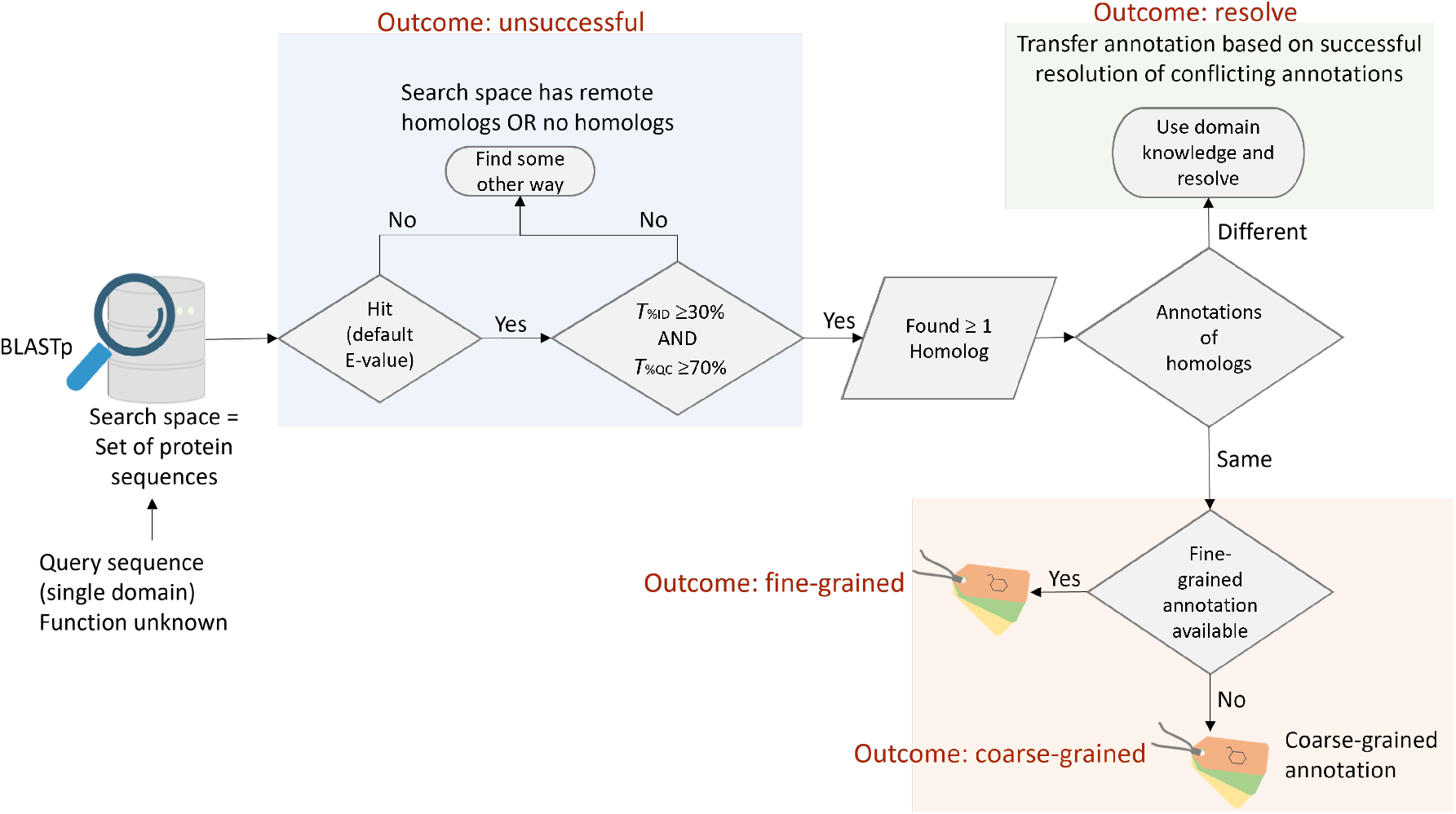
Flowchart of possible outcomes in fine-grained molecular function annotation. The figure illustrates the decision flow for assigning fine-grained molecular function to a protein using BLASTp against a curated functional database. Three possible outcomes are shown: (i) No homologs found - molecular function remains unassigned, though remote homologs may exist (unsuccessful). (ii) Homologs found with consistent annotations - annotation transfer depends on the granularity of the function (fine-grained or coarse-grained). (iii) Homologs with conflicting annotations - domain knowledge is needed to resolve conflicts, depending on ground truth data availability (resolve).

### Multiple sequence alignment

Algorithms for multiple sequence alignments (MSA) assume that all the input sequences are homologs, i.e., they have evolved from a common ancestor. A column in an MSA therefore shows the residues in the input sequences that map to the same amino acid position in the ancestral sequence [19]. However, it becomes increasingly difficult to identify (align) homologous positions in input sequences as the level of sequence divergence increases. This is particularly so if the number of indel events is high. Unfortunately, there is no mathematical or statistical parameter (something similar to the e-value that BLAST uses) that one can use to assess the quality of MSAs. A set of benchmarks, such as HOMSTRAD and/or BAliBASE databases, is typically used to compare the performance of various MSA algorithms.

The accuracy and cost of nine popular MSA programs, namely CLUSTALW, CLUSTAL OMEGA, DIALIGN-TX, MAFFT, MUSCLE, POA, PROBALIGN, PROBCONS, and T-COFFEE, were compared against the benchmark alignment dataset BAliBASE [20]. The accuracy of alignment was calculated as the sum-of-pairs and total-column scores. The computational costs were determined by collecting peak memory usage and time taken for the execution of the program. This comparative study found that consistency-based programs, viz., PROBCONS, T-COFFEE, PROBALIGN, and MAFFT, outperformed the rest. It further found that MAFFT can deliver faster and reliable alignments. In another study [21], the authors highlighted that MAFFT can consistently produce correct alignments with high accuracy in comparison to the other five MSA programs, viz., CLUSTALW, DIALIGN-T, MUSCLE, PROBCONS, and T-COFFEE. Hence, MAFFT v7.490(2012/Oct/30), installed on a Linux server, was used to obtain MSAs. The mafft command was used to generate an alignment file (.aln) for the given protein family. This alignment file was visually analyzed using Jalview software [22] and further examined for gaps and conservation. Default values were used for all the parameters.

### Hidden Markov models

We utilized profile hidden Markov models (HMMs) [23], [11], [24] to determine if a given protein sequence belongs to the protein family represented by the profile HMM. HMMER 3.3 software package was installed on a Linux server. The workflow has four steps:

i. A multiple sequence alignment (MSA) of the set of homologous sequences belonging to a protein family was generated using the MAFFT alignment tool [25].
ii. A separate profile HMM was built for each protein family from the MSA using the hmmbuild command.
iii. The bit score threshold for each profile HMM was set by first performing self-scoring, i.e., scoring every sequence that was used to build the profile HMM against the same profile HMM. This gave a range of best 1 domain bit scores (Tables 1 and 2 in Supporting information S1 File), and these ranges were used as guides to plot ROC curves (Figs 2-6 in Supporting information S1 File). Bit score thresholds (*T*_ROC_) were set such that there are no false positives when the search space is the collection of sequences from all the datasets used in the present study (Supporting information S1 Table). We note that *T*_ROC_ is dependent on the search space.
iv. Every query sequence was scored against the profile HMM using the `hmmsearch` command. If the bit score of a query sequence was *≥ T*_ROC_ of the protein family’s profile HMM, the sequence was inferred to belong to that family. In other words, the query sequence was considered homologous to the sequences in the protein family.

We used the `best 1 domain` bit score for further analysis for the following reasons: (i) Unlike E-value, bit score does not depend on the size of the search space. Hence, we get the same value whether we score a sequence by itself or as a part of a collection of sequences. (ii) The best 1 domain score contains the bit score for the single best scoring domain, unlike the score which sums up scores for all (partial) alignments as well as sequences repeats, if any, present in a sequence (https://eddylab.org/software/hmmer/Userguide.pdf).

**Fig 2.**
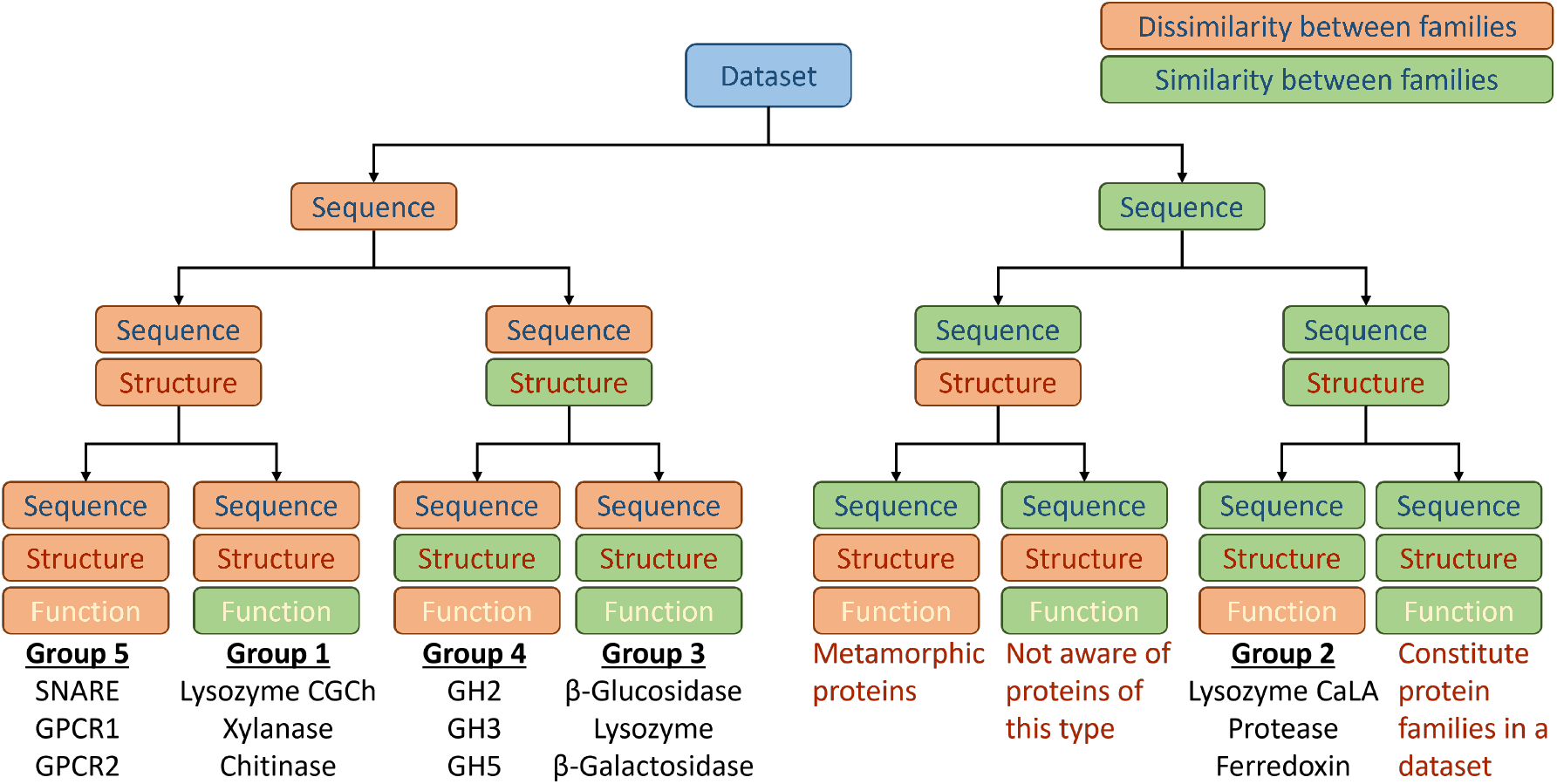
Sequence-structure-function relationships in protein families across datasets. The protein families included in the datasets exhibit different combinations of sequence, structure, and function similarity, which can be either similar or dissimilar. These datasets are organized into five groups, covering all combinations except three. The missing combinations are: (i) sequence similarity without structural and functional similarity, typically observed in metamorphic proteins (inadequate data) [32], [33]; (ii) sequence and functional similarity without structural similarity, a relationship which we are not aware (inadequate data); and (iii) similarity in sequence, structure, and function, which defines proteins belonging to the same family in this study.

## Results

We present the results as follows: (A) Detection of homology of every sequence with other sequences (i) within the family, (ii) of protein families within the dataset, and (iii) from other families considered in this study. (B) To assess whether annotations transferred based on inference are correct. We have chosen proteins with known functions so that we can find out if transferred annotations are correct or not. Based on such an assessment, we discuss the suitability of BLASTp and profile HMM when the query sequences are of unknown function (fine-grained).

### Using BLASTp to assess if sequences considered in a protein family are homologs

Datasets were created in such a way that proteins within each protein family are homologous to each other, as inferred from their shared fold. However, the percentage of alignment pairs exceeding the BLASTp threshold for homology within a family (%*H*_intra_) varies from 22.6% (Cellulase; Group 4) to 100.0% (Lysozyme G; Group 1) (see %*H*_intra_ in Tables 4 and 5). This variation is largely due to the sequence divergence caused by the broad taxonomic distribution leading to remote homology. For example, the Cellulase family in the GH5 dataset (Group 4) includes sequences from Eukaryota, Bacteria, and Archaea; some sequences are Unclassified. GH10 family in Xylanase dataset (Group 1) displays moderate conservation (%*H*_intra_ = 52.1%), includes Eukaryota and Bacteria. In contrast, the sequences from the Lysozyme G family of Lysozyme CGCh (Group 1) are all from Eukaryota. These observations are consistent with the well-documented limitation of BLASTp to detect remote homology.

**Table 4.**
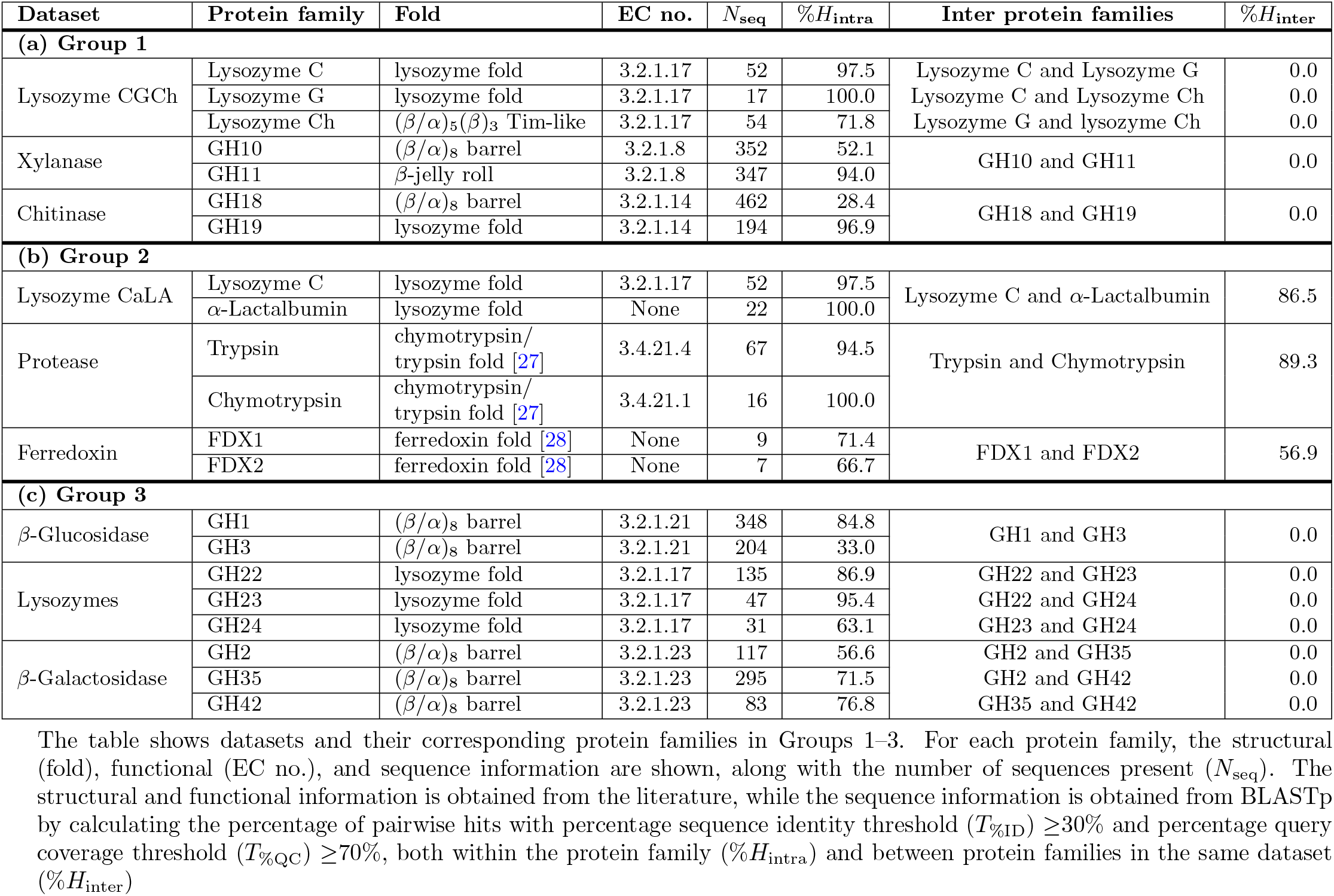
Structural, functional, and sequence information of protein families in Groups 1–3.

**Table 5.**
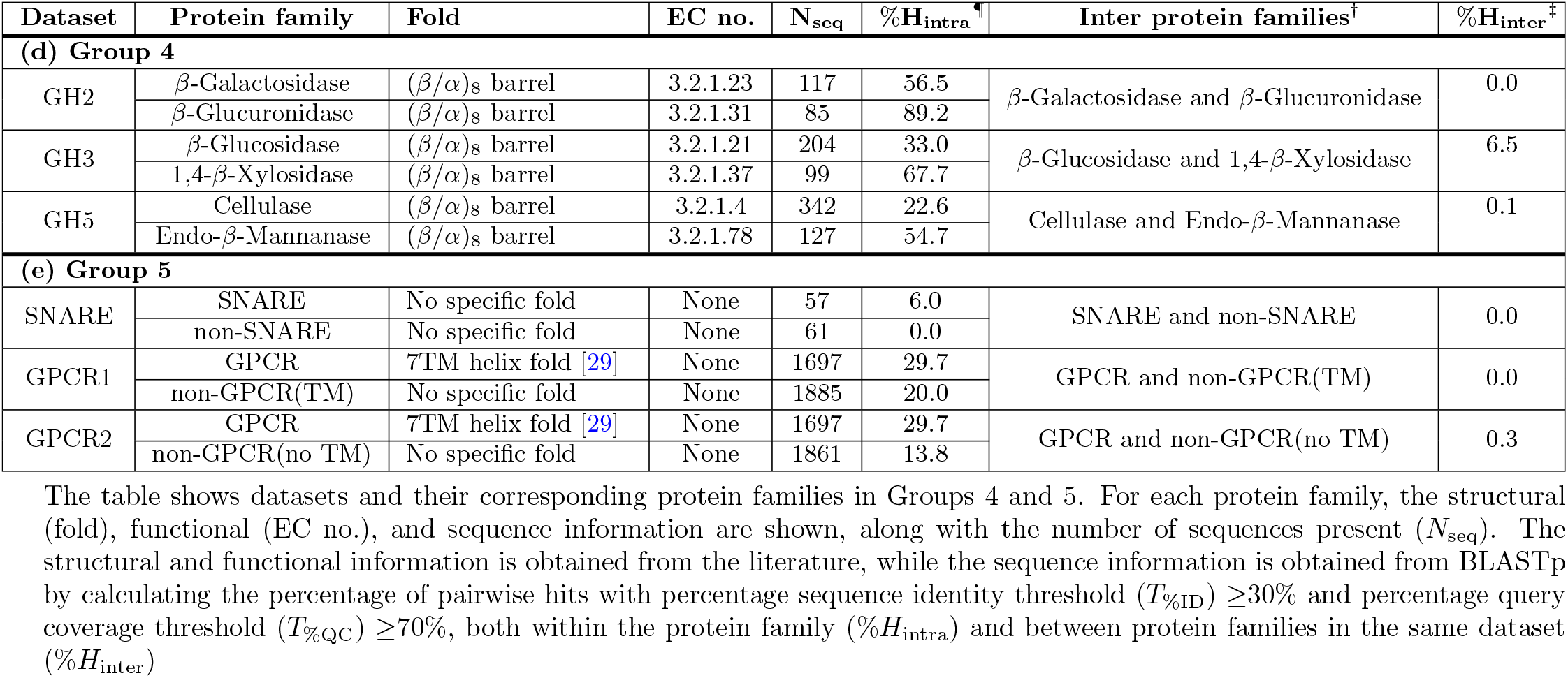
Structural, functional, and sequence information of protein families in Groups 4 and 5.

### Using BLASTp to assess if protein families of a dataset are homologs

The protein families in the datasets of Groups 2, 3, and 4 are homologous to each other due to the same fold (Table 1). BLASTp is able to detect homology in Group 2 (%*H*_inter_ ranges from 56.9% to 89.3%, Tables 4 and 5) but not in Group 3 and Group 4 (%*H*_inter_ either 0.0% or close to 0.0%). This is because of the inability of BLASTp to detect remote homologs caused by sequence divergence. Sequence divergence can be due to broader taxonomic distribution of sequences as well as the differences in rate of evolution of different protein families (section 3.3c in [26]). For example, in Group 2, the protein sequences in all datasets were from Eukaryota, whereas those in Group 3 and Group 4 were from Archaea, Bacteria, Eukaryota, and Viruses (only in Group 3; Lysozyme dataset), along with some unclassified.

Protein families in Group 1 datasets are non-homologous by design (Table 1). BLASTp not detecting any homologs is as expected. Even in Group 5, the choice of proteins is such that there is no homology between SNARE and non-SNARE (as described in the Methods section); so is the case with GPCR and non-GPCR (TM)/non-GPCR (no TM). Hence, %*H*_inter_ = 0.0% as expected (see %*H*_inter_ in Table 5).

### Alignment of every sequence with every other sequence considered in this study using BLASTp

So far, we have analyzed results in the specific context of sequence identity within and between the protein families of a dataset. We also analysed results when sequences were aligned with sequences belonging to other families considered in this study. BLASTp finds homology of several sequences from Groups 1-4 with sequences belonging to the non-SNARE and non-GPCR sequences of Group 5 (Supporting information S2 Table). These are not unexpected because non-SNARE and non-GPCR sequences were chosen with the only condition that they are not SNAREs/GPCRs (with or without TM domain) as described in the method section.

### Assessment of annotations based on homology inferred from BLASTp alignments

We present the assessment using the workflow shown in Fig 1. The assessment considers a search space which is the collection of sequences from all the datasets used in the present study. This assessment is applicable as is even when faced with a protein of unknown function, its sequence is typically used as a query in a BLASTp search against a reference database. This database can be a curated collection (e.g., SwissProt, CAZy, UniRef, or a user-defined set) or a large aggregated resource containing all known sequences (e.g., NCBI nr).

Consider a protein for which we are seeking to assign fine-grained molecular function. The sequence of this protein is used as the query for BLASTp search against a database consisting of proteins with curated molecular function. We can envisage the possible outcomes (Fig 1): (i) No homologs are found. One may choose other search spaces to find homologs. Assuming that the search spaces represent all known molecular functions, BLASTp is of no help in assigning molecular function. We may note that the presence of remote homologs cannot be ruled out (Outcome: unsuccessful). (ii) Homologs are found with consistent molecular function. In this case, transfer of annotation will be justified, and the granularity of function annotation is determined by annotations of homologs (Outcome: fine-grained or Outcome: coarse-grained). (iii) Annotations of the homologs are not identical. Here, one needs to use domain knowledge to resolve the conflicting annotation (Outcome: resolve) but its success depends on the nature of the available ground truth data.

Herein, we analyse the outcome of the above workflow for each sequence when queried against the search space comprising all other sequences considered in this study (Supporting information S2 Table). We also provide the detailed annotations of subject sequences that are different from the query sequence. These details are taken from the UniProt. Analysis of query sequences whose hits do not have the same annotations (Outcome: resolve) is given below, with additional details for this analysis given in Supporting information S3 Table.

i. Lysozyme C, *α*-lactalbumin, and sperm acrosome membrane-associated protein 3 (Spaca3): these are sequence and structural homologs that perform completely different functions. Resolution by a domain expert depends upon the level of sequence identity of the query with these three functionally different groups of proteins and whether active site residues are conserved.
ii. Lysozyme C and GH22 family proteins: All these proteins catalyse the same reaction (same at the fine-grained level) but are involved in different biological processes. At the coarse-grained level, they are the same with respect to the reaction catalysed (glycoside hydrolase) but different with respect to the biological process.
iii. GH19 family chitinase and endochitinase 1: Both are glycoside hydrolases (same at coarse-grained level) but differ in their reaction specificity.
iv. Trypsin, chymotrypsin, and serine protease/peptidase: All these proteins catalyse the same reaction, hydrolysis of the peptide bond (same in coarse-grained level), but differ from each other in terms of their cleavage site and biological processes.
v. FDX1 and FDX2: These two isoforms are highly specific for their substrates and participate in distinct biochemical pathways. The distinct conserved sequence motifs should be used to annotate FDX1 or FDX2 while resolving the conflicting annotation [30], [28].
vi. GH1 family *β*-glucosidase and myrosinase 2: The former is a *β*-glucosidase, whereas the latter is a thioglucosidase. Whether each of the two enzymes can also act on the substrates of the other enzyme is not known. Hence, the available data is inadequate to resolve.
vii. GH3 family *β*-glucosidases and 1,4-*β*-xylosidases: *β*-glucosidases are exo-enzymes and some are known to have broader substrate specificity, i.e., hydrolyse *β*-D-galactosides, *α*-L-arabinosides, *β*-D-xylosides, and *β*-D-fucosides also (same at coarse-grained level). Hence, an unambiguous resolution of conflicting annotations is not possible.
viii. Cellulase and endo-*β*-mannanases: Both are glycoside hydrolases (same at coarse-grained level) but differ in substrate specificity. Hence, an unambiguous resolution of conflicting annotations is not possible.

### Using profile HMMs to assess if sequences considered in a protein family are homologs

Sequences belonging to each protein family were scored against the corresponding profile HMM (Table 6). The number of sequences whose bit score against the profile HMM is lower than *T*_ROC_ even though these sequences were used to build the profile HMM, is denoted as *N*_low_. All the sequences of the family meet the bit score threshold in nearly half of the cases, i.e., *N*_low_ = 0. For other families *N*_low_≠ 0 because our aim is to choose *T*_ROC_ such as to make FPR = 0 i.e., avoid propagation of wrong annotation even if it means the bit score for some of the sequences of the family <*T*_ROC_.

**Table 6.**
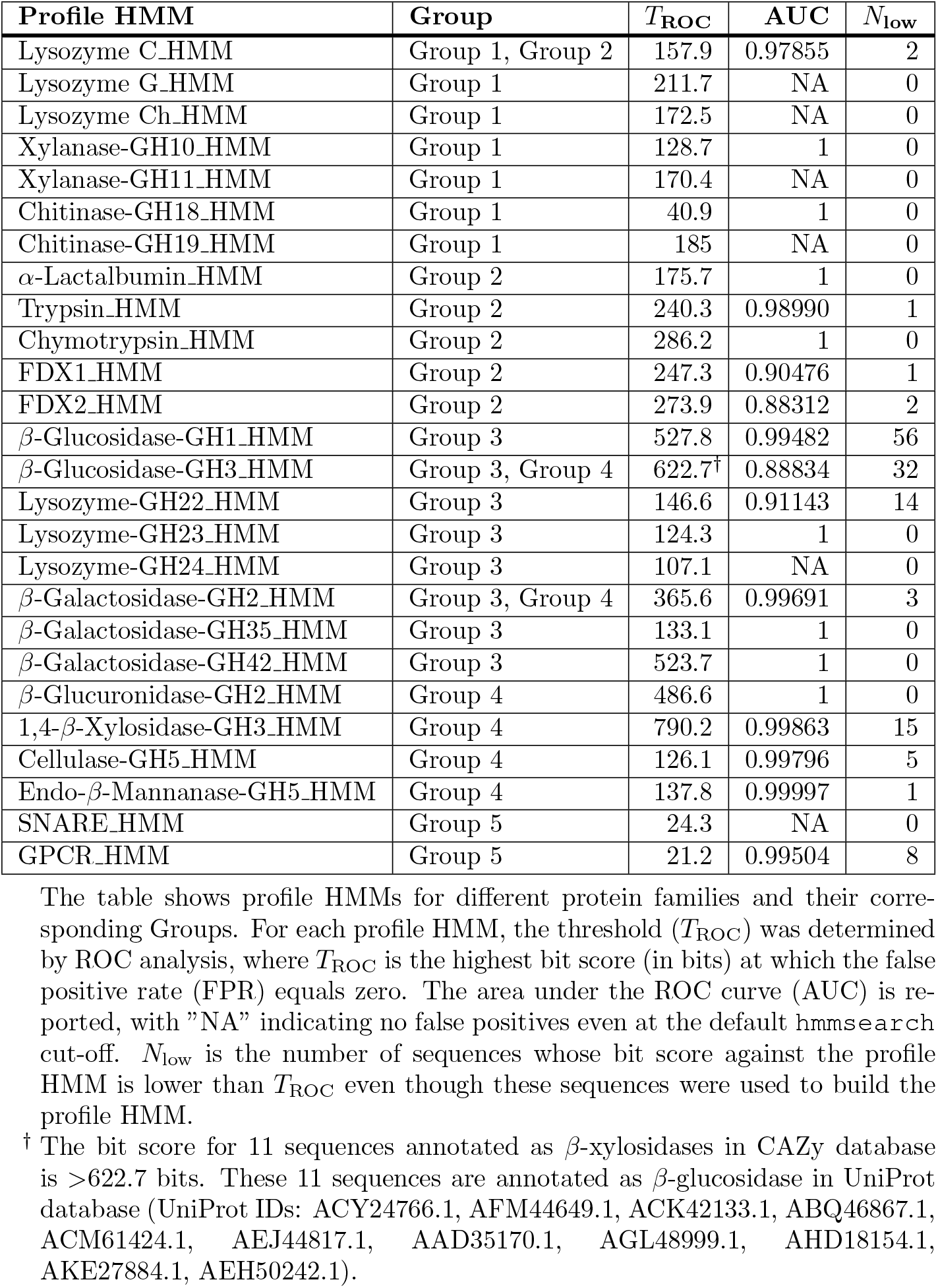
Profile HMM thresholds and performance on protein families of all Groups.

| Profile HMM | Group | $T_{\text{ROC}}$ | AUC | $N_{\text{low}}$ |
| --- | --- | --- | --- | --- |
| Lysozyme C_HMM | Group 1, Group 2 | 157.9 | 0.97855 | 2 |
| Lysozyme G_HMM | Group 1 | 211.7 | NA | 0 |
| Lysozyme Ch_HMM | Group 1 | 172.5 | NA | 0 |
| Xylanase-GH10_HMM | Group 1 | 128.7 | 1 | 0 |
| Xylanase-GH11_HMM | Group 1 | 170.4 | NA | 0 |
| Chitinase-GH18_HMM | Group 1 | 40.9 | 1 | 0 |
| Chitinase-GH19_HMM | Group 1 | 185 | NA | 0 |
| α-Lactalbumin_HMM | Group 2 | 175.7 | 1 | 0 |
| Trypsin_HMM | Group 2 | 240.3 | 0.98990 | 1 |
| Chymotrypsin_HMM | Group 2 | 286.2 | 1 | 0 |
| FDX1_HMM | Group 2 | 247.3 | 0.90476 | 1 |
| FDX2_HMM | Group 2 | 273.9 | 0.88312 | 2 |
| β-Glucosidase-GH1_HMM | Group 3 | 527.8 | 0.99482 | 56 |
| β-Glucosidase-GH3_HMM | Group 3, Group 4 | 622.7<sup>†</sup> | 0.88834 | 32 |
| Lysozyme-GH22_HMM | Group 3 | 146.6 | 0.91143 | 14 |
| Lysozyme-GH23_HMM | Group 3 | 124.3 | 1 | 0 |
| Lysozyme-GH24_HMM | Group 3 | 107.1 | NA | 0 |
| β-Galactosidase-GH2_HMM | Group 3, Group 4 | 365.6 | 0.99691 | 3 |
| β-Galactosidase-GH35_HMM | Group 3 | 133.1 | 1 | 0 |
| β-Galactosidase-GH42_HMM | Group 3 | 523.7 | 1 | 0 |
| β-Glucuronidase-GH2_HMM | Group 4 | 486.6 | 1 | 0 |
| 1,4-β-Xylosidase-GH3_HMM | Group 4 | 790.2 | 0.99863 | 15 |
| Cellulase-GH5_HMM | Group 4 | 126.1 | 0.99796 | 5 |
| Endo-β-Mannanase-GH5_HMM | Group 4 | 137.8 | 0.99997 | 1 |
| SNARE_HMM | Group 5 | 24.3 | NA | 0 |
| GPCR_HMM | Group 5 | 21.2 | 0.99504 | 8 |

The table shows profile HMMs for different protein families and their corresponding Groups. For each profile HMM, the threshold ( $T_{\text{ROC}}$ ) was determined by ROC analysis, where $T_{\text{ROC}}$ is the highest bit score (in bits) at which the false positive rate (FPR) equals zero. The area under the ROC curve (AUC) is reported, with "NA" indicating no false positives even at the default hmmsearch cut-off. $N_{\text{low}}$ is the number of sequences whose bit score against the profile HMM is lower than $T_{\text{ROC}}$ even though these sequences were used to build the profile HMM.
<sup>†</sup> The bit score for 11 sequences annotated as $\beta$ -xylosidases in CAZy database is >622.7 bits. These 11 sequences are annotated as $\beta$ -glucosidase in UniProt database (UniProt IDs: ACY24766.1, AFM44649.1, ACK42133.1, ABQ46867.1, ACM61424.1, AEJ44817.1, AAD35170.1, AGL48999.1, AHD18154.1, AKE27884.1, AEH50242.1).

A possible reason as to why *N*_low_ ≠ 0 for *β*-Glucosidase-GH3 HMM and 1,4-*β*-Xylosidase-GH3 HMM is the similarity shared by their substrates at the monomeric level: the substituent at C-5 is -CH_2_OH in glucopyranose and -H in xylopyranose. Several *β*-xylosidases score higher than *β*-glucosidases against the *β*-Glucosidase-GH3 HMM and vice versa for the 1,4-*β*-Xylosidase-GH3 HMM. In the case of *β*-Glucosidase-GH1 HMM, *N*_low_ = 56 because myrosinase 2 (UniProt ID: Q9C5C2), a thioglucosidase (EC 3.2.1.147) from Non-GPCR (no TM) set, scores 527.2 bits score against the *β*-Glucosidase-GH1 HMM.

The *N*_low_ ≠ 0 for Lysozyme-GH22 HMM because of two possible reasons: (i) sperm acrosome membrane-associated protein 3 (UniProt: Q9D9X8) scores *≥ T*_ROC_. It is a sequence and structural homolog of lysozyme C which has lost enzymatic activity because of the mutation of catalytic site residues [31]. (ii) 13 of the 135 sequences of this family score much below the *T*_ROC_ because of high sequence divergence relative to the other 122 sequences.

### Using profile HMMs to assess homology across protein families and datasets

While using BLASTp, thresholds for *T*_%ID_ and *T*_%QC_ to infer homology between the query and subject sequences are independent of the sequences being aligned. In contrast, each profile HMM will have a customized bit score threshold. As mentioned in the Methods section, bit score thresholds were set such that there are no false positives when the search space is the collection of sequences from all the datasets used in the present study. In view of this, none of the sequences score higher than *T*_ROC_ for profile HMMs of other families. For example, if we consider the Protease dataset, none of the chymotrypsins are hits for Trypsin HMM and vice versa. Because of these reasons, setting a bit score threshold carefully can eliminate the possibility of erroneous transfer of function annotation.

The workflow for searching a database using a profile HMM for the purpose of transferring annotation (Fig 7 in Supporting information S1 File) is broadly similar to the one described for BLASTp (Fig 1) except that the we use bit score threshold (*T*_ROC_) instead of percentage sequence identity threshold (*T*_%ID_) and percentage query coverage threshold (*T*_%QC_).

## Discussion and Conclusion

High-throughput genome sequencing has led to the accumulation of a very large number of conceptually translated protein sequences. Knowledge of the function(s) performed by such proteins is essential to fully realise the value of high-throughput sequencing projects. It is neither practical to experimentally determine the function(s) of such a large number of proteins nor desirable. Nevertheless, the sequence data can be effectively used to assign molecular functions to proteins of interest, and this can be experimentally validated. BLASTp and profile-based methods (PSI-BLAST, profile HMM) are widely used to assign function based on homology. However, assignment of molecular function based on homology may be erroneous because these methods do not by default distinguish orthologs from paralogs. Hence, in this study, we have investigated the suitability or effectiveness of BLASTp and profile HMM to assign fine-grained molecular function based on inferred homology using carefully curated datasets that cover diverse sequence-structurefunction relationships (Table 1) (Fig 2).

BLASTp thresholds (*T*_%ID_ and *T*_%QC_) to infer homology between query and subject sequences are inde-pendent of the sequences being aligned. Because of this reason we do find homologs which do not share function, for example, in Group 2 datasets. Transfer of function annotation in these cases will obviously be erroneous. In Group 4 datasets, homology cannot be inferred because of high sequence divergence, and hence the question of transferring annotation does not arise.

In contrast to BLASTp, profile HMMs have customized bit score thresholds for inferring homologs. A sequence that scores above this threshold is inferred as belonging to the protein family represented by the profile HMM. Thresholds may be stringent so as to make the profile ’specific’ (eliminate false positives) or relaxed so that the profile is sensitive (eliminate false negatives). In this study, we chose to prioritize specificity over sensitivity. It is possible to adopt a two-step workflow wherein a relaxed threshold is used in the first step so that there are no false negatives. In the second step, hits from the first step that do not meet the stringent threshold may be taken up for further analysis.

One may choose to generate separate profiles for each paralogous family to eliminate the possibility of erroneous transfer of function as illustrated by the Protease dataset: none of the chymotrypsin sequences are hits for the Trypsin HMM and vice versa.

We have presented a general workflow to annotate protein sequences using BLASTp or profile HMM against any search space. In our study, a collection of all sequences except the one being used as the query constituted the search space. The question of transferring annotation does not arise if the query does not have any homologs in the search space. If homologs are found, then the decision to transfer annotation is straightforward if all hits have the same annotation. This is not the case when hits with different annotations are obtained; differences which may be due to (i) mutation at the active site leading to loss or change of function (e.g., lysozyme C, *α*-lactalbumin, and Spaca3), (ii) inadequate experimental data leading to ambiguous annotation (e.g., GH1 family *β*-glucosidase and myrosinase), and (iii) function being same at coarse-grained level but different at fine-grained level (e.g., trypsin and chymotrypsin) due to differences in substrate utilized, reaction catalysed, and/or biological process. Such conflicting annotations have to be resolved by domain experts. However, this may or may not be possible as illustrated below.

The case of Spaca3 meeting the threshold of Lysozyme-GH22 HMM is easier to resolve because of the mutation of active site residues in Spaca3. Checking for the conservation of active site residues will suffice to infer whether hits of this Lysozyme-GH22 HMM are Spaca3. *β*-glucosidase and 1,4-*β*-xylosidase satisfy the thresholds of both *β*-Glucosidase-GH3 HMM and 1,4-*β*-Xylosidase-GH3 HMM. Activities of all *β*-glucosidases against xylosides have not been tested; similarly, activities of all *β*-xylosidases have not been tested against glucosides. It may turn out that *β*-glucosidase and 1,4-*β*-xylosidase are orthologs with broad substrate specificity but these are considered as paralogs. The current knowledge of the sequencesubstrate specificity relationship is inadequate to resolve whether these enzymes are paralogs or orthologs without experimental data. Another such example is *β*-glucosidase of *β*-Glucosidase-GH1 HMM which acts on O-glucosides, and myrosinase 2 which acts on thioglucosides. In general, enzymes are known to display promiscuity with respect to substrate utilized, reaction catalysed, and/or product formed [34], [35]. Often, data on promiscuity is not available because of the additional resources required to test for promiscuity. Paucity of such data limits the use of profile HMMs for transferring annotation.

Several proteins are known to have moonlighting function (http://www.moonlightingproteins.org/) [36]. Moonlighting, unlike promiscuity, does not affect bit score thresholds. One can envisage two possibilities: within a family of homologous proteins, some but not all have moonlighting function. This leads to uncertainties about transferring annotations related to additional (moonlighting) functions. We also note that just because a protein can perform a function does not necessarily mean it does so in vivo, as shown for *SipB* [37]. This illustrates that although transfer of function annotation based on inferred homology seems to be justified, the protein may not carry out the annotated function in vivo. Addressing challenges such as these requires experimental data.

In conclusion, sequence divergence, variations in protein lengths, and annotation granularity within a protein family mean that considering a larger number of proteins for generating profile HMMs makes the profile less reliable if the interest is in transferring function at the fine-grained level. Erroneous function transfer may occur, especially at the fine-grained level due to limited experimental data, subtle sequence-function relationships leading to substrate, reaction, and product promiscuities, and biological ingenuities such as moonlighting proteins. Nevertheless, the present study demonstrates the use of BLASTp and profile HMMs to annotate protein functions while highlighting a set of challenging cases that preclude automation.

## Acknowledgment

We thank the Indian Institute of Technology Bombay for the infrastructure and other facilities.

## Supporting information

- **S1 File**
- **S1 Table**
- **S2 Table**
- **S3 Table**

## Footnotes

[1 some researchers set the sequence identity threshold to 25% [8], [9]

