## Supplementary material for "Systematic assessment of homology-based methods for fine-grained functional annotation using diverse protein families": S1 File

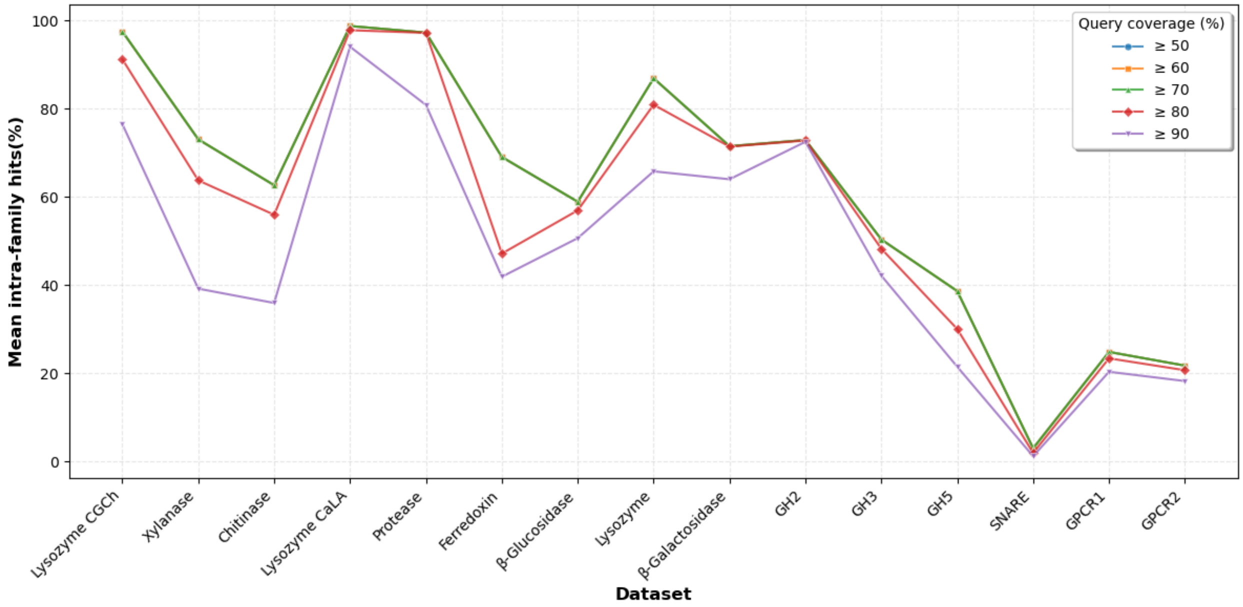


**Fig 1. Comparison of $T_{\%QC}$ thresholds on intra-family BLASTp hits across datasets.** The threshold of percentage query coverage ($T_{\%QC}$) for inferring homology based on pairwise BLASTp at threshold of percentage sequence identity ($T_{\%ID}$) $\geq$30% was evaluated using intra-family hits (%), since sequences within protein families are homologs. We calculated the mean intra-family hits (%) for all protein families in each dataset under varying $T_{\%QC}$ thresholds. The percentage of homologs detected by setting $T_{\%QC}$ threshold to $\geq$50%, $\geq$60%, or $\geq$70% remains the same. Making the threshold more stringent ($\geq$80% or $\geq$90%) expectedly reduces the percentage of homologs detected. In view of this, $T_{\%QC}$ was set to $\geq$70% in our study.

**Table 1. Domain-bit score ranges of profile HMM searches within datasets of Groups 1-4.**

| Dataset | Protein family | Profile HMM | | |
|---|---|---|---|---|
| **(a) Group 1** | | | | |
| | | Lysozyme G_HMM | Lysozyme C_HMM | Lysozyme Ch_HMM |
| Lysozyme CGCh | Lysozyme G | 360.7 - 243.4 | 0.0 - 0.0 | 0.0 - 0.0 |
| | Lysozyme C | 0.0 - 0.0 | 253.9 - 146.5 | 0.0 - 0.0 |
| | Lysozyme Ch | 0.0 - 0.0 | 0.0 - 0.0 | 370.5 - 172.5 |
| Xylanase | | Xylanase-GH10_HMM | Xylanase-GH11_HMM | |
| | GH10 | 487.4 - 128.7 | 0.0 - 0.0 | |
| | GH11 | 0.0 - 0.0 | 406.7 - 170.4 | |
| Chitinase | | Chitinase-GH18_HMM | Chitinase-GH19_HMM | |
| | GH18 | 507.0 - 40.9 | 12.1 - 12.1 | |
| | GH19 | 0.0 - 0.0 | 531.2 - 185.0 | |
| **(b) Group 2** | | | | |
| Lysozyme CaLA | | Lysozyme C_HMM | $\alpha$-Lactalbumin_HMM | |
| | Lysozyme C | 253.9 - 146.5 | 138.3 - 68.7 | |
| | $\alpha$-Lactalbumin | 145.0 - 95.6 | 267.0 - 175.7 | |
| Protease | | Trypsin_HMM | Chymotrypsin_HMM | |
| | Trypsin | 366.2 – 212.6 | 252.8 – 116.0 | |
| | Chymotrypsin | 231.2 - 186.4 | 421.9 – 286.2 | |
| Ferredoxin | | FDX1_HMM | FDX2_HMM | |
| | FDX1 | 295.4 - 117.9 | 205.6 - 73.4 | |
| | FDX2 | 167.9 - 72.3 | 310.8 - 90.0 | |
| **(c) Group 3** | | | | |
| $\beta$-Glucosidase | | BGL-GH1_HMM | BGL-GH3_HMM | |
| | GH1 | 660.4 - 174.6 | 0.0 - 0.0 | |
| | GH3 | 0.0 - 0.0 | 988.1 - 246.5 | |
| Lysozyme | | LYZ-GH22_HMM | LYZ-GH23_HMM | LYZ-GH24_HMM |
| | GH22 | 217.1 - 37.1 | 0.0 - 0.0 | 0.0 - 0.0 |
| | GH23 | 0.0 - 0.0 | 324.7 - 124.3 | 0.0 - 0.0 |
| | GH24 | 0.0 - 0.0 | 0.0 - 0.0 | 209.4 - 107.1 |
| $\beta$-Galactosidase | | $\beta$-Gal-GH2_HMM | $\beta$-Gal-GH35_HMM | $\beta$-Gal-GH42_HMM |
| | GH2 | 1400.9 - 300.7 | 16.9 - 10.3 | 0.0 - 0.0 |
| | GH35 | 16.8 - 9.3 | 1148.0 - 133.1 | 34.2 - 8.0 |
| | GH42 | 25.3 - 8.1 | 34.5 - 8.7 | 1001.8 - 523.7 |
| **(d) Group 4** | | | | |
| GH2 | | $\beta$-Gal-GH2_HMM | GUS-GH2_HMM | |
| | $\beta$-Galactosidase | 1400.9 - 300.7 | 268.8 - 99.6 | |
| | $\beta$-Glucuronidase | 334.0 - 242.0 | 931.3 - 486.6 | |
| GH3 | | BGL-GH3_HMM | BXL-GH3_HMM | |
| | $\beta$-Glucosidase | 988.1 - 246.5 | 779.0 - 130.8 | |
| | 1,4-$\beta$-Xylosidase | 809.8 - 215.4 | 1037.5 - 575.1 | |
| GH5 | | Cellulase-GH5_HMM | MAN-GH5_HMM | |
| | Cellulase | 338.8 - 55.0 | 137.2 - 18.5 | |
| | Endo-$\beta$-Mannanase | 124.3 - 26.0 | 451.7 - 104.1 | |

The table shows the `best 1 domain` bit score (DBS) range obtained by searching profile HMMs of protein families against all protein families within the same dataset, including self-search. The analysis includes datasets from Groups 1–4. The following abbreviations are used for profile HMM names: (i) BGL-GH1_HMM: $\beta$-Glucosidase-GH1_HMM, (ii) LYZ-GH22_HMM: Lysozyme-GH22_HMM, (iii) $\beta$-Gal-GH2_HMM: $\beta$-Galactosidase-GH2_HMM, (iv) GUS-GH2_HMM: $\beta$-Glucuronidase-GH2_HMM, (v) BXL-GH3_HMM: 1,4-$\beta$-Xylosidase-GH3_HMM, and (vi) MAN-GH5_HMM: Endo-$\beta$-Mannanase-GH5_HMM. Note: DBS after the inclusion threshold was not included.

**Table 2. Domain-bit score ranges of profile HMM searches within datasets of Group 5.**

| **(a) Group 5** | | |
|---|---|---|
| **Dataset** | **Protein family** | **Profile HMM** |
| | | SNARE_HMM |
| SNARE | SNARE | 132.1 - 24.3 |
| | non-SNARE | 0.0 - 0.0 |
| GPCR1 | | GPCR_HMM |
| | GPCR | 287.8 - 8.8 |
| | non-GPCR (TM) | 35.2 - 20.3 |
| GPCR2 | | GPCR_HMM |
| | GPCR | 287.8 - 8.8 |
| | non-GPCR (no TM) | 0.0 - 0.0 |

The table shows the `best 1 domain` bit score (DBS) range obtained by searching profile HMMs of protein families against all protein families within the same dataset, including self-search. The analysis includes datasets from Group 5. Note: DBS after the inclusion threshold was not included.

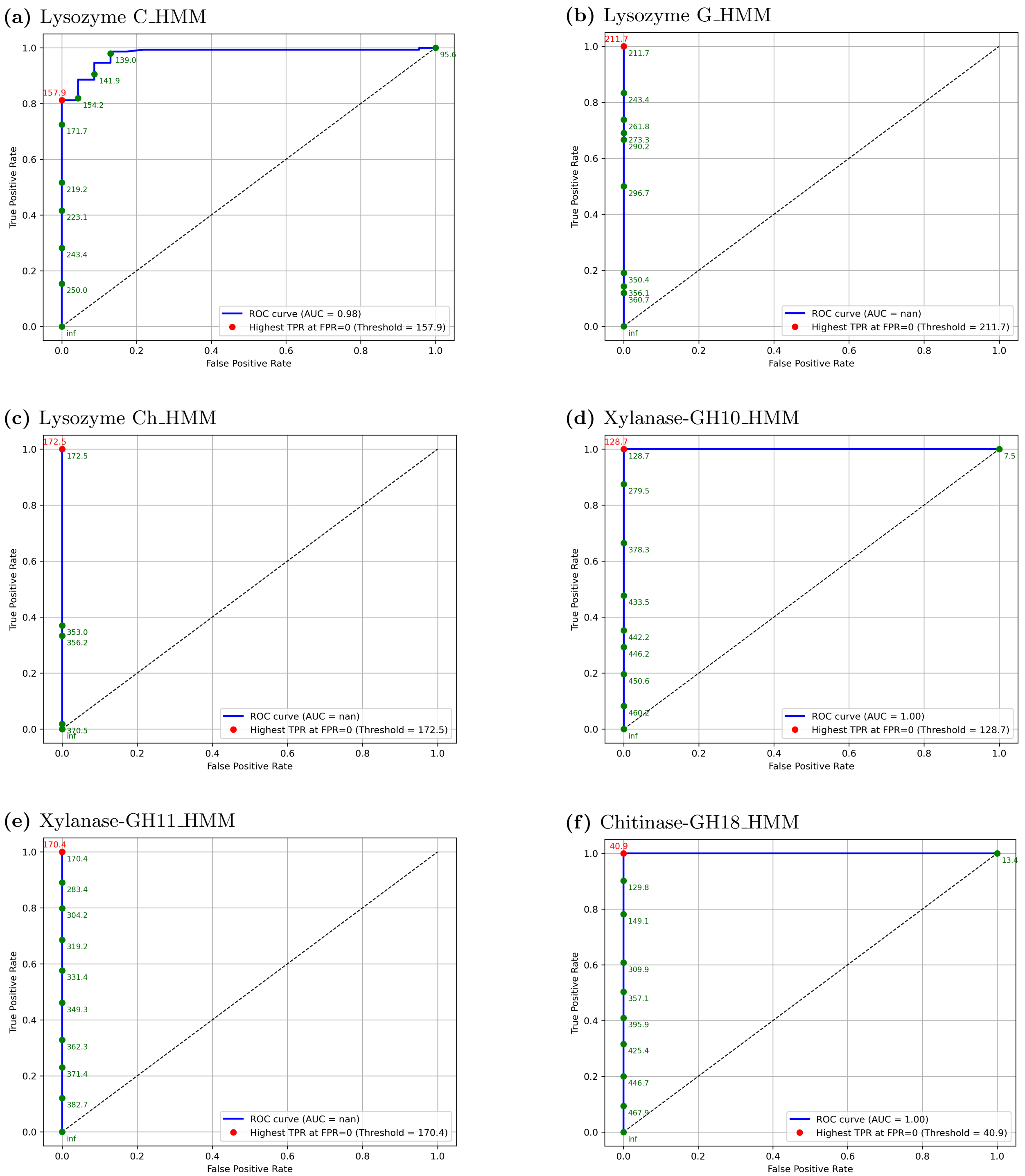


**Fig 2. AUC ROC curves for six profile HMMs.** Each figure shows the AUC ROC curve corresponding to one of six profile HMMs of different protein families tested across all datasets. The area under the curve (AUC) was calculated for each ROC curve, and the threshold ($T_{\mathrm{ROC}}$) - defined as the point with the highest true positive rate (TPR) at false positive rate (FPR) = 0, is highlighted in red. An AUC value of "nan" indicates that no false positives were detected, even using the default `hmmsearch` parameters.

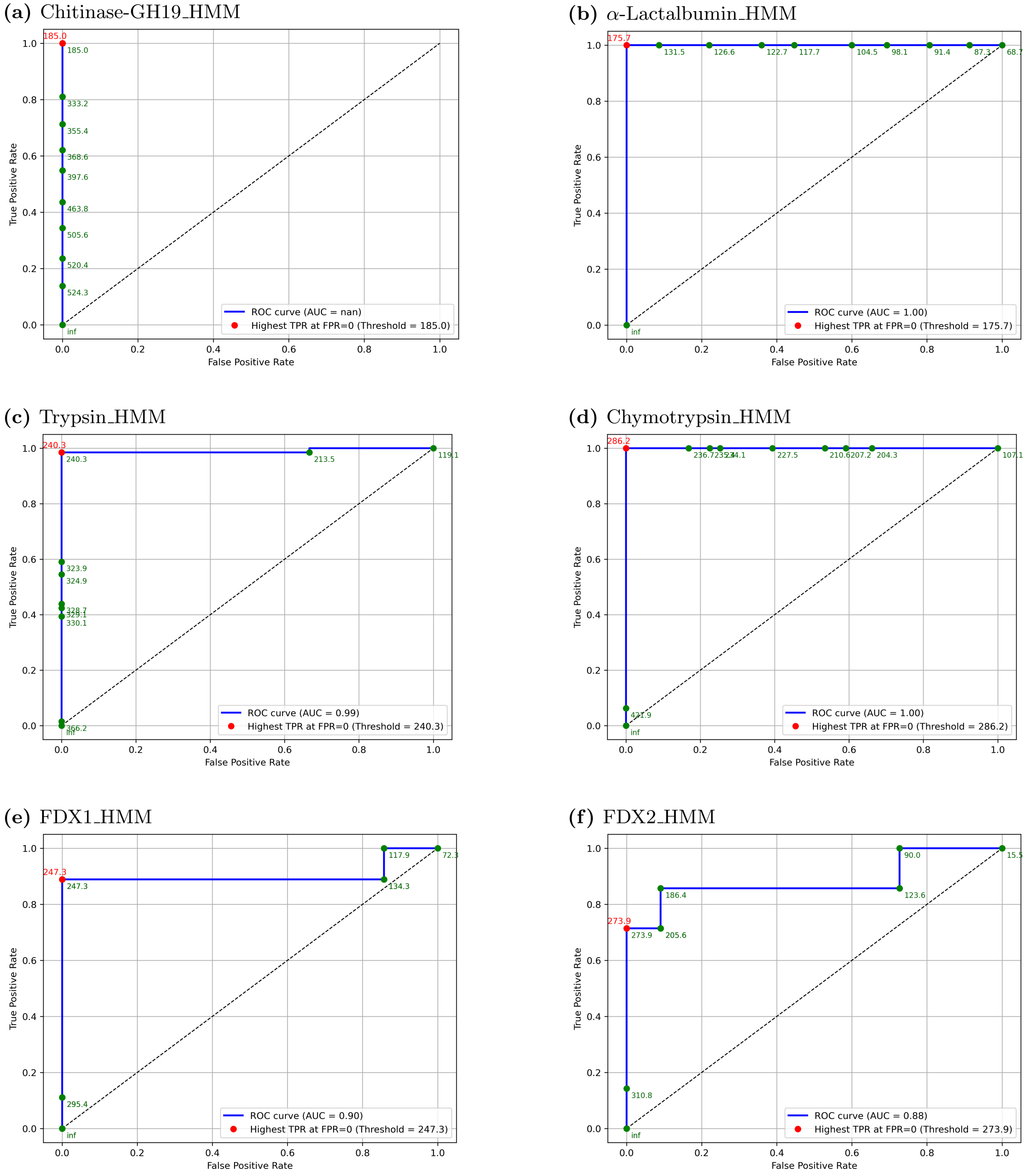


**Fig 3. AUC ROC curves for six profile HMMs.** Each figure shows the AUC ROC curve corresponding to one of six profile HMMs of different protein families tested across all datasets. The area under the curve (AUC) was calculated for each ROC curve, and the threshold ($T_{\mathrm{ROC}}$) - defined as the point with the highest true positive rate (TPR) at false positive rate (FPR) = 0, is highlighted in red. An AUC value of "nan" indicates that no false positives were detected, even using the default `hmmsearch` parameters.

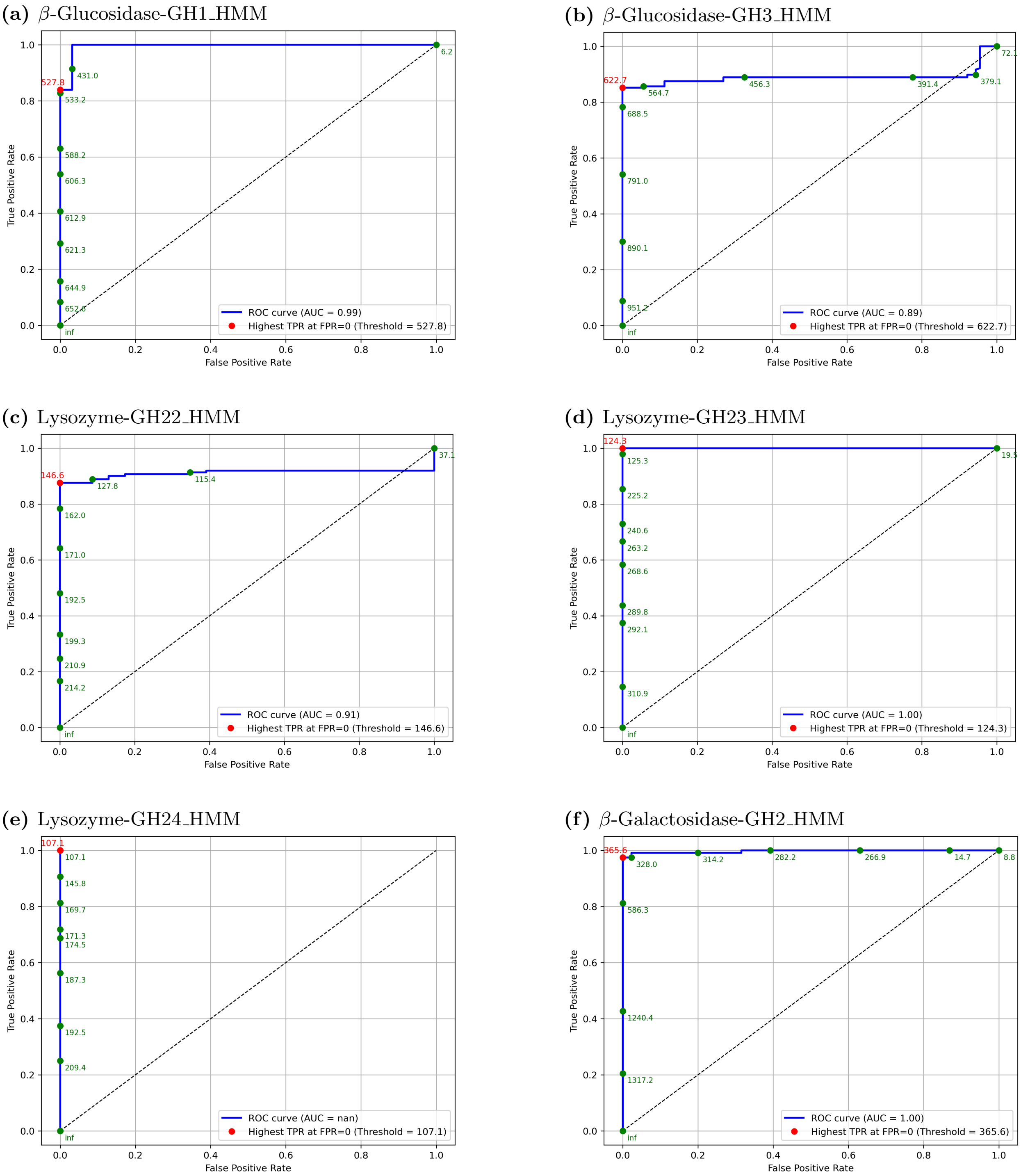


**Fig 4. AUC ROC curves for six profile HMMs.** Each figure shows the AUC ROC curve corresponding to one of six profile HMMs of different protein families tested across all datasets. The area under the curve (AUC) was calculated for each ROC curve, and the threshold ($T_{\mathrm{ROC}}$) - defined as the point with the highest true positive rate (TPR) at false positive rate (FPR) = 0, is highlighted in red. An AUC value of "nan" indicates that no false positives were detected, even using the default `hmmsearch` parameters.

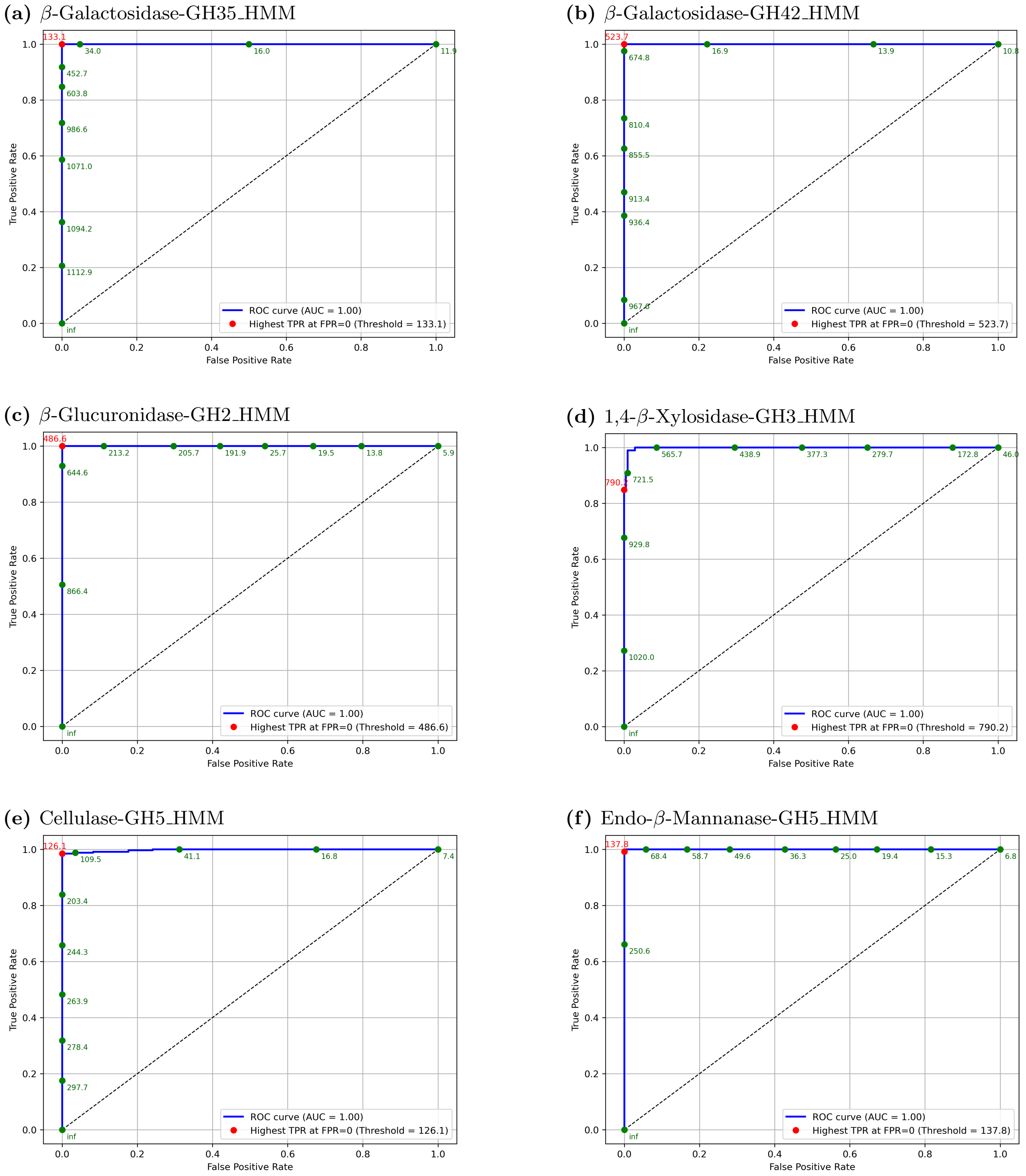


**Fig 5. AUC ROC curves for six profile HMMs.** Each figure shows the AUC ROC curve corresponding to one of six profile HMMs of different protein families tested across all datasets. The area under the curve (AUC) was calculated for each ROC curve, and the threshold ($T_{\mathrm{ROC}}$) - defined as the point with the highest true positive rate (TPR) at false positive rate (FPR) = 0, is highlighted in red.

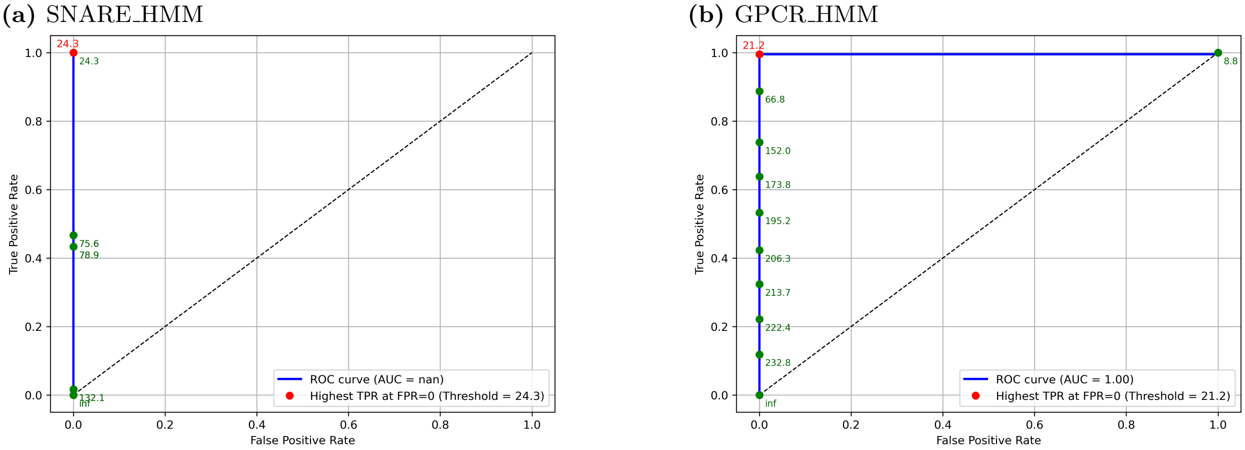


**Fig 6. AUC ROC curves for two profile HMMs.** Each figure shows the AUC ROC curve corresponding to one of two profile HMMs of different protein families tested across all datasets. The area under the curve (AUC) was calculated for each ROC curve, and the threshold ($T_{\mathrm{ROC}}$) - defined as the point with the highest true positive rate (TPR) at false positive rate (FPR) = 0, is highlighted in red. An AUC value of "nan" indicates that no false positives were detected, even using the default `hmmsearch` parameters.

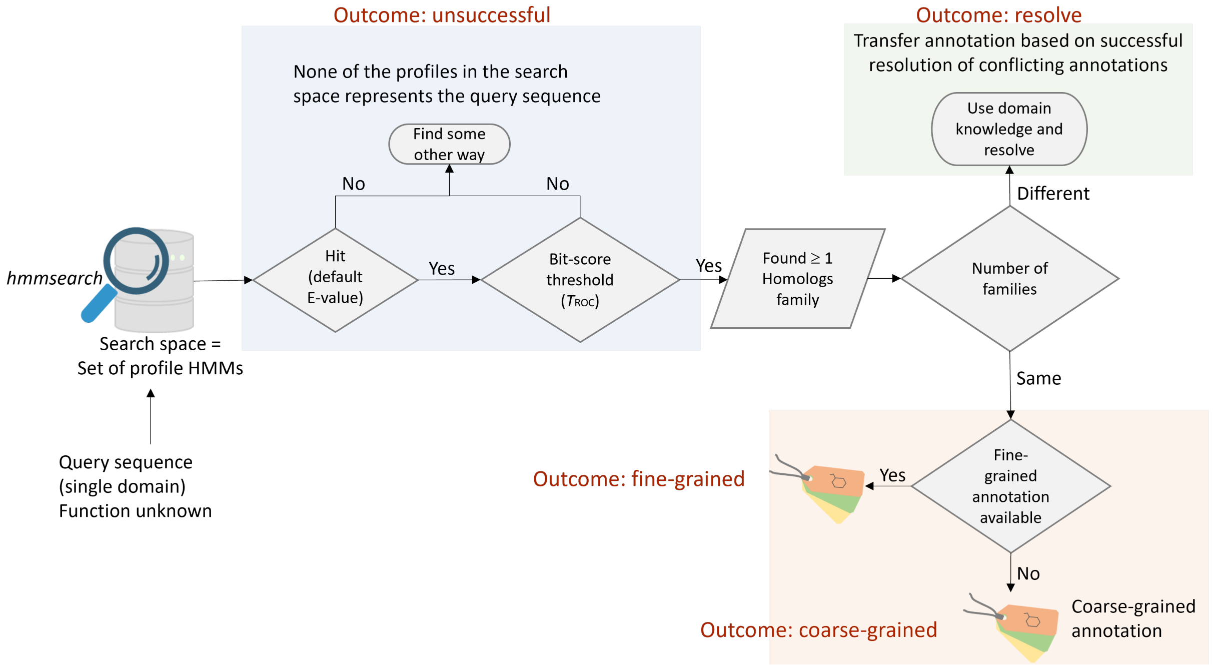


**Fig 7. Flowchart of possible outcomes in fine-grained molecular function annotation.** The figure illustrates the decision flow for assigning fine-grained molecular function to a protein using profile HMM against a curated functional database. Three possible outcomes are shown: (i) No homologs found - molecular function remains unassigned, though remote homologs may exist (unsuccessful). (ii) Homologs found with consistent annotations - annotation transfer depends on the granularity of the function (fine-grained or coarse-grained). (iii) Homologs with conflicting annotations - domain knowledge is needed to resolve conflicts, depending on ground truth data availability (resolve).
